# Structured nucleosome fingerprints enable high-resolution mapping of chromatin architecture within regulatory regions

**DOI:** 10.1101/016642

**Authors:** Alicia N. Schep, Jason D. Buenrostro, Sarah K. Denny, Katja Schwartz, Gavin Sherlock, William J. Greenleaf

## Abstract

Transcription factors canonically bind nucleosome-free DNA, making the positioning of nucleosomes within regulatory regions crucial to the regulation of gene expression. We observe a highly structured pattern of DNA fragment lengths and positions generated by the assay of transposase accessible chromatin (ATAC-seq) around nucleosomes in *S. cerevisiae*, and use this distinctive two-dimensional nucleosomal “fingerprint” as the basis for a new nucleosome-positioning algorithm called NucleoATAC. We show that NucleoATAC can identify the rotational and translational positions of nucleosomes with up to base pair resolution and provide quantitative measures of nucleosome occupancy in *S. cerevisiae*, *S. pombe*, and human cells. We demonstrate application of NucleoATAC to a number of outstanding problems in chromatin biology, including analysis of sequence features underlying nucleosome positioning, promoter chromatin architecture across species, identification of transient changes in nucleosome occupancy and positioning during a dynamic cellular response, and integrated analysis of nucleosome occupancy and transcription factor binding.

## Main Text

Chromatin accessibility modulates the ability of transcription factors (TFs) and transcriptional machinery to interact with DNA. Within regions of increased accessibility, nucleosomes and TFs compete for access to regulatory DNA1. While sequence content has been shown to influence nucleosome positioning, the specific locations of nucleosomes *in vivo* are also dynamically modulated by chromatin remodelers, transcription factors, and transcriptional machinery^2–4^. Therefore, methods for producing base-pair resolved nucleosome maps with quantitative occupancy information within regulatory elements promise to provide insight into the interplay between chromatin organization and transcriptional regulators—a crucial step toward a comprehensive and predictive understanding of how regulatory elements control gene expression.

Methods for inferring nucleosome positions through MNase digestion of chromatin followed by high-throughput sequencing of protected fragments have provided a window into the relationship between TF binding and nucleosome positioning, revealing that certain TFs are consistently flanked by well-positioned nucleosomes while others show considerable heterogeneity in the positioning of proximal nucleosomes^4–6^. However, MNase-based methods are limited in their ability to infer high-resolution nucleosome positions and provide quantitative measures of nucleosome occupancy by the enzyme’s processive nature of DNA digestion and intrinsic digestion sequence bias^7,8^. Alternatively, chemical mapping approaches in both *Saccharomyces cerevisiae* and *Schizosaccharomyces pombe* have provided base-pair resolved maps of nucleosome positions^9,10^. However, this high-resolution chemical cleavage technique cannot be easily adapted to other biological systems, as it requires a genetically modified histone H4. Furthermore, as with MNase-based assays, chemical mapping has limited ability to measure absolute nucleosome occupancy, as nucleosome depletion is indirectly inferred through lack of signal.

We recently described the assay for transposase-accessible chromatin using sequencing (ATAC-seq), a method for rapid, sensitive genome-wide, profiling of chromatin accessibility^11^. Here we adapt ATAC-seq to *S. cerevisiae* and discover a highly structured, reproducible ATAC-seq fragmentation pattern around nucleosomes. We use this “nucleosome fingerprint” as the basis of NucleoATAC, a computational method for quantitative, high-resolution inference of nucleosome positioning and occupancy within regulatory regions. We show that NucleoATAC infers nucleosome positions at near bp-resolution in *S. cerevisiae,* demonstrated by high concordance with chemical mapping data, and captures rotational positioning of nucleosomes. Furthermore, we show that the stereotyped “nucleosome fingerprint” derived from *S. cerevisiae* can be applied to data sets from different species, including humans, thereby generating the first high-resolution nucleosome calls in human regulatory regions. We highlight several applications of NucleoATAC by examining differences in chromatin architecture in regulatory regions between the three species examined, elucidating changes in nucleosome positioning and occupancy during a dynamic transcriptional response in yeast, and determining nucleosome occupancy and positioning relative to transcription factors in a human lymphoblastoid cell line.

## Results

### V-plots demonstrate structured ATAC-seq signal around nucleosomes

We previously observed that short ATAC-seq fragments are concentrated at nucleosome-free regions, whereas long fragments are enriched at nucleosome-associated DNA^11^. To further examine this association, we developed an *S. cerevisiae* ATAC-seq protocol to determine ATAC-seq fragmentation patterns at positions of base-pair resolved nucleosomes in *S. cerevisiae* generated using chemical mapping techniques^9^. Using ATAC-seq for *S. cerevisiae* we generated 61 million paired-end ATAC-seq reads with high mapping quality across 11 replicates (**Supplementary Fig. 1**; Methods). ATAC-seq read depth for *S. cerevisiae* is highly correlated with DNase-seq^12^ (**Fig. 1a; Supplementary Fig. 2a**) but shows greater enrichment in promoters (**Supplementary Fig. 2b**), demonstrating that ATAC-seq provides a sensitive measure of chromatin accessibility genome-wide. As with mammalian ATAC-seq, fragment sizes for *S. cerevisiae* reflect nucleosome organization, with a peak in the fragment size distribution at 140-200 bps arising from DNA protected by a nucleosome (**Fig. 1b**).

**Figure 1.**
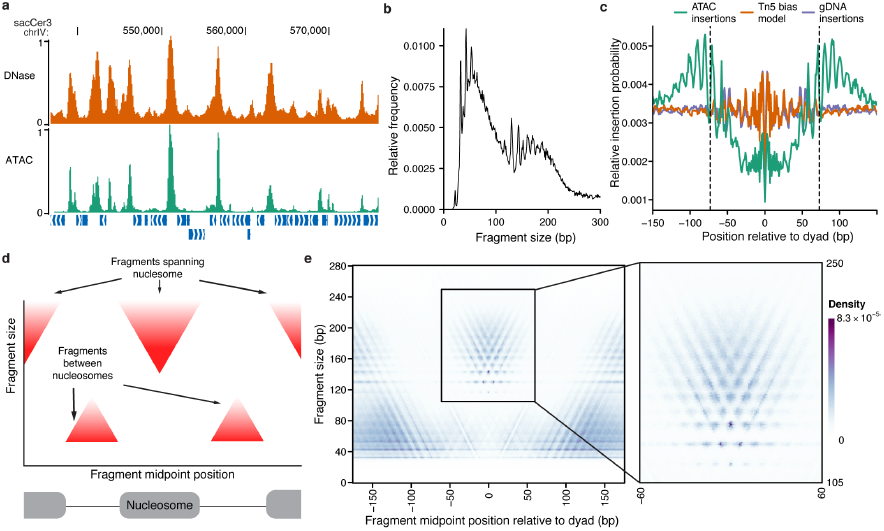
ATAC-seq signal is highly structured around nucleosomes. A) ATAC-seq (green) insertion track for *S. cerevisiae* shows enrichment of insertions at accessible chromatin regions, similar to DNase-seq cut density (orange). Both tracks were smoothed by 150 bp and scaled so that the maximum density in the region is 1. B) Fragment size distribution for *S. cerevisiae* ATAC-seq samples. C) Insertion probabilities for ATAC-seq (teal), genomic DNA (purple) and predicted by sequence bias (orange; see methods) around nucleosomes defined by chemical mapping. D) Schematic illustration of expected V-plot pattern around a well-positioned nucleosome E) V-plot (fragment size versus fragment center position) of ATAC-Seq fragments around well-positioned nucleosomes called by chemical mapping, with inset showing region with nucleosome-spanning fragments.

By aggregating ATAC-seq transposition centers around well-positioned, base-pair resolved nucleosome positions determined by chemical mapping^9^, we observe clear protection from transposase insertion within nucleosomal DNA (**Fig. 1c**). Additionally, we observe striking periodicity in the insertions at the boundary of the nucleosome. We postulate that this periodicity arises from steric hindrance of the Tn5 transposase at the nucleosome boundary, which allows for only one face of the DNA double helix to be accessible to transposition. Using the nucleosome crystal structure^13^, we observe the insertion hotspots at 58 and 71 bps from the dyad correspond to sites where the major groove of DNA is facing away from the nucleosome (**Supplementary Fig. 3**).

To further characterize the ATAC-seq signal around these nucleosome dyad positions, we mapped fragment midpoints and sizes using a “V-plot”^14^ (**Fig. 1d**). This visualization maps the density of fragment sizes versus fragment center locations relative to a genomic feature of interest (in this case, nucleosome dyads). These aggregate protection profiles show a V-shaped structure, where the apex of the “V” represents the smallest possible fragment that spans the DNA protected by a nucleosome. The V-plot centered on chemically mapped dyads shows a clear depletion of short fragments in the portion of DNA wrapped around the nucleosome (**Fig. 1e**). At fragment sizes spanning a nucleosome (**Fig. 1e inset**), we observe a highly structured V-pattern with both horizontal and vertical periodicity. This periodicity likely reflects both the steric hindrance of the transposase (vertical and horizontal periodicity) and previously described 10 bp rotational positioning of nucleosomes in yeast (horizontal periodicity). The apex of the V shape is at 117 bp while the most abundant position in the V-plot represents fragments of 143 bps centered at the dyad. These smaller than expected fragment sizes may arise from stochastic “breathing” of DNA associated with nucleosomes allowing for transposase insertions within the 147 bp that are canonically considered to be nucleosome associated^15^ or from nucleosomes packed closer than 147 bp apart^16^.

### Determining nucleosome positions from structured V-plot

We reasoned that standard methods for inferring nucleosome centers, which assume that fragment mid-points are normally distributed around the nucleosome core^17,18^, could be improved by leveraging this highly structured two dimensional V-plot pattern. To this end, we developed NucleoATAC (**Fig. 2**), an algorithm that cross-correlates the characteristic, average nucleosome V-plot against a V-plot representation of fragments across the genome (see Methods). This cross-correlation signal measures how well ATAC-seq data at any particular base fits the expected pattern at a nucleosome dyad. To account for the possibility of spurious signal from Tn5 insertion sequence bias^11,19^ and signal variation based on differential chromatin openness, we normalize this nucleosome signal by subtracting a calculated background signal expected from transposition sequence bias, the global fragment size distribution of the sample, and the number of fragments in the region. Peaks from the background-subtracted signal track are used to identify dyad positions, which are then scored for several characteristics that can be used for downstream filtering (see Methods). To determine nucleosome occupancy, the global fragment size distribution is modeled as a mixture of two distributions, nucleosomal and nucleosome-free (**Supplementary Fig. 4**), and the maximum likelihood fraction of nucleosomal reads at a locus is taken as the occupancy score.

**Figure 2.**
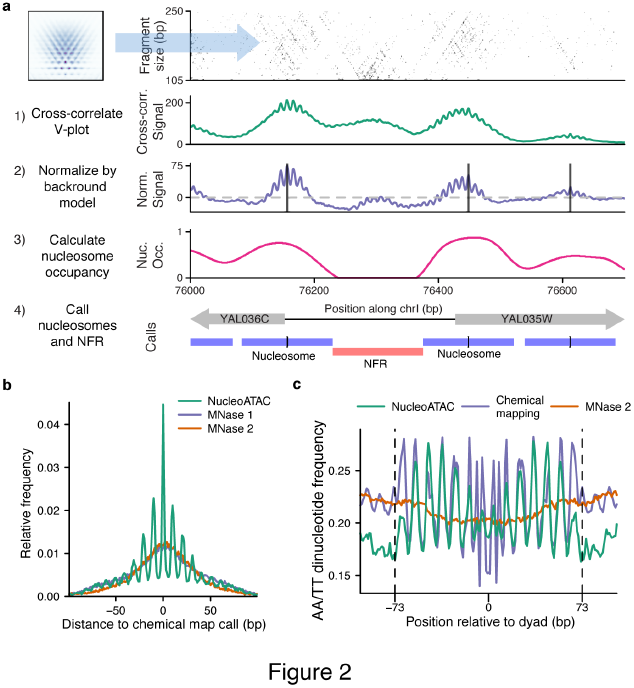
NucleoATAC enables high-resolution nucleosome positioning. A) Schematic of NucleoATAC workflow. First, the V-plot nucleosome signature is cross-correlated against a 2D fragment size versus fragment midpoint representation of ATAC-seq data at a locus. The signal is then normalized by a background model (based on sequence bias and read depth) to obtain a normalized signal. Nucleosome occupancy is calculated using the local fraction of nucleosomal fragments. The normalized cross-correlation signal and nucleosome occupancy tracks are used to assign nucleosome and nucleosome-free (NFR) positions. B) Distance of NucleoATAC dyad calls (green) and two sets of MNase dyad calls (purple and orange) from chemical mapping dyad calls. C) AA/TT dinucleotide pattern around nucleosome dyad calls determined by NucleoATAC (green), chemical mapping (purple), and MNase (orange).

### NucleoATAC enables high-resolution nucleosome calling in *S. cerevisiae*

NucleoATAC identified the positions of 40,049 nucleosomes across the yeast genome (with Z-score >= 3, log-likelihood ratio > 0, occupancy > 0.1; see Methods). We found that no characteristic nucleosome fingerprint is observed when aggregating ATAC-seq insertions generated from genomic DNA or predicted by transposase bias signal (**Supplementary Fig. 5**) around NucleoATAC-called nucleosomes, suggesting residual intrinsic Tn5 insertion bias has little effect on aggregate NucleoATAC nucleosome calls.

Nucleosome positioning calls determined by NucleoATAC are highly concordant (**Fig. 2b**) with chemically mapped nucleosomes^9^. Divergent calls are generally offset by multiples of 10 base-pairs, demonstrating that NucleoATAC captures the 10 bp rotational positioning of nucleosomes, unlike calls from MNase^20,21^ datasets (**Fig. 2b**). This rotational positioning of nucleosome calls enables us to observe the underlying sequence periodicity that may dictate the rotational positioning of nucleosomes; AA/TT dinucleotide content exhibits strong periodicity within DNA contacting NucleoATAC nucleosomes **(Fig. 2c)**. The confidence metrics used for filtering NucleoATAC calls enable the algorithm to be robust to sequencing depth; when down-sampling or using individual replicates from our data, fewer nucleosomes are called but called nucleosomes have similar concordance with the chemical mapping calls (**Supplementary Fig. 6, 7**). As positional concordance between NucleoATAC and chemical mapping calls increases as a function of both NucleoATAC and chemical map confidence metrics (**Supplementary Fig. 8**), discrepancies between the two methods are likely partially due to either lower quality chemical mapping calls or inconsistently positioned nucleosomes.

### NucleoATAC can be applied across species

Because histones are among the most evolutionarily conserved proteins, we hypothesized that the same structured V-plot “nucleosome fingerprint” pattern is present for different species. To test this possibility, we first developed ATAC-seq for *Schizosaccharomyces pombe*, a species for which high-resolution chemical mapping data are also available^10^. Subtle aspects of the ATAC-seq fragment size distribution for *S. pombe* differed from *S. cerevisiae*, as might be expected based on previously characterized differences in average linker lengths^10^. However, local maxima in the distributions aligned relatively well **(Fig. 3a)**, suggesting similar nucleosomal constraints on insertion between the species. We reasoned that adjusting the *S. cerevisiae* V-plot so that the relative intensities of the rows would match the nucleosomal insert size distribution of the *S. pombe* sample (see Methods) would approximate the *S. pombe* nucleosome V-plot. Indeed, the resulting V-plot is similar to a V-plot made from ATAC-seq reads using *S. pombe* chemically mapped dyads (R = 0.96 after normalization; R = 0.81 without normalization) (**Fig. 3b,c**). We applied NucleoATAC using both the adjusted *S. cerevisiae* V-plot and the “true” V-plot from *S. pombe* chemical map calls, and found the resulting calls to be very similar (**Supplementary Fig. 9**) and highly concordant with the chemical mapping calls **(Fig. 4d)**. These results suggest that nucleosomal constraints on the Tn5 transposase are conserved across species and that NucleoATAC can be applied to any organism of interest.

**Figure 3.**
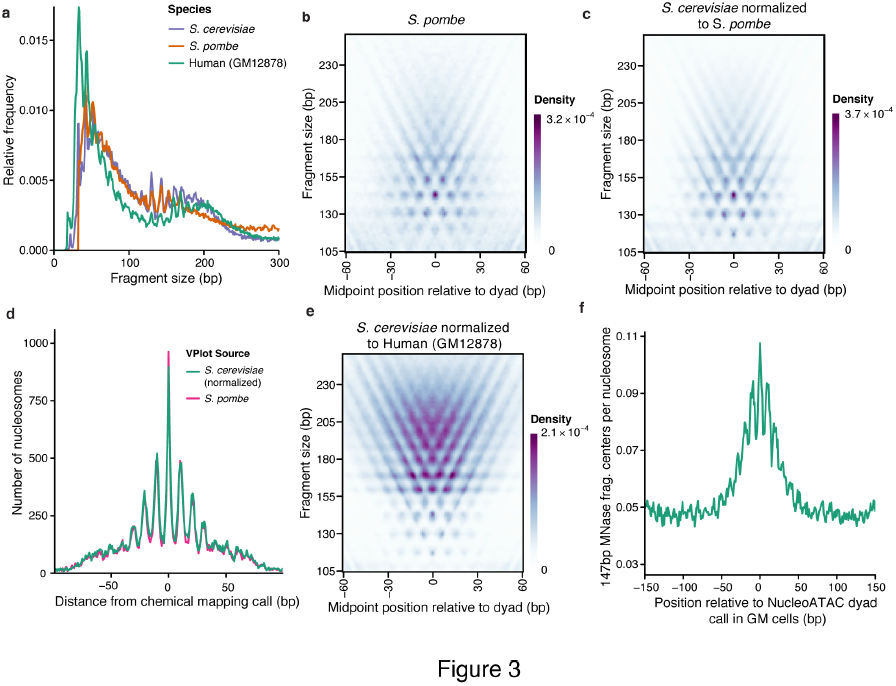
V-plot derived from *S. cerevisiae* can be used as a template to apply NucleoATAC to other species. A) Fragment sizes distributions for *S. cerevisiae* (purple), *S. pombe* (orange), and human GM12878 cell line (teal) B) *S. pombe* V-plot based on chemical map calls for *S. pombe* C) *S. cerevisiae* V-plot normalized to match *S. pombe* fragment size distribution D) Comparison of NucleoATAC concordance with chemical mapping for *S. pombe* when using V-plots in (B) or (C). E) *S. cerevisiae* V-plot normalized to match human GM12878 fragment sizes F) 147 bp MNase fragment density around calls for GM12878 made by NucleoATAC.

**Figure 4.**
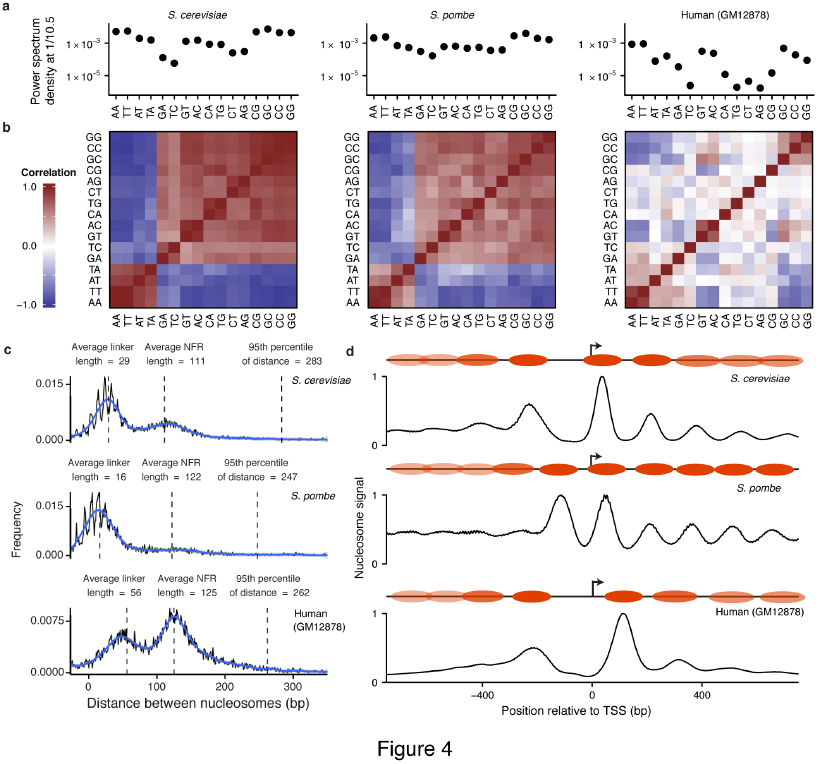
NucleoATAC reveals differences in nucleosome architecture between species. A) Power spectrum density at 1/10.5 bp for each dinucleotide from 19 to 60 bp from NucleoATAC-called dyads for *S. cerevisiae*, *S. pombe*, and human (left to right). B) Pair-wise correlation between dinucleotide frequencies for each species. C) Distances between adjacent nucleosomes (with low intervening nucleosome occupancy) in three species. Annotation shows typical linker and NFR lengths, as well as the 95^th^ percentile in distance. D) Positive NucleoATAC signal aggregated at TSS in three species. Cartoons show canonical nucleosome positioning at TSS for each species, with more transparent nucleosome ovals representing nucleosomes that are less consistently positioned among different TSSs.

To apply NucleoATAC to ATAC-seq data from the human lymphoblastoid GM12878 cell line^11^, we similarly normalized the V-plot signal density from *S. cerevisiae* to match the nucleosomal fragment size distribution of the human data **(Fig. 3e)**. Unlike yeast, much of the human epigenome is heterochromatic and therefore yields low ATAC-seq signal; thus analysis was restricted to broad open chromatin peaks called by MACS2^22^. To assess our ability to capture rotational positioning of human nucleosomes, we compared our calls with the 147 bp fragment center positions from MNase-seq experiments – fragments that have been shown to display 10 bp periodicity in dinucleotide content^5^. The positions of these 147 bp MNase fragment centers display clear 10 bp periodicity around NucleoATAC calls (**Fig. 3f**), validating our ability to capture rotational information in human cells.

### Comparison of nucleosome positioning across species

High-resolution NucleoATAC nucleosome calls and signal tracks allowed for a comprehensive and quantitative analysis of sequence preferences of well-positioned nucleosomes in regulatory regions across these species. Dinucleotide frequencies between 19 to 60 bps from the nucleosome dyad often displayed a 10.5 bp periodicity, tracking the helical pitch of DNA around the nucleosome (**Supplementary Fig. 10**). We computed a pair-wise correlation of this enrichment between every dinucleotide for each species (**Fig. 4a**), as well as the power spectrum density at frequency 1/10.5 bp for each dinucleotide as a measure of the magnitude of this oscillation (**Fig. 4b**). For the two yeast species, all dinucleotides show strong 10.5 bp periodicity, with the AA, TA, AT, and TT dinucleotides being out of phase with all the other dinucleotides. For human nucleosomes, we observe overall much weaker periodicity, with the strongest relative periodicity in AA, TT, GT, AC, GC, CC, and GG, and only weak or negligible power for other di-nucleotides. The weaker periodicity in CG relative to GC, GG, and CC might reflect the effect of CpG methylation, in line with other work suggesting that methylated CpG frequencies oscillate out of phase with un-methylated CG^23^.

Our high-resolution calls allow for a detailed comparison of regulatory architecture across these three disparate species. For each species, the distribution of distances between the boundaries of nucleosomes (with low intervening occupancy scores) is bimodal, with one peak representing typical linker length and the other nucleosome-free region (NFR) length. Linker length varies between the species, with *S. pombe* having the shortest linker length and human the longest, consistent with previous observations^10,24^. For both yeast species we observe “negative” linker lengths representing two nucleosomes with dyads being closer than 147 bp (**Fig. 4c**), supporting findings from paired-end chemical mapping and the hypothesis that chromatin can exist in a state with partially unwrapped nucleosomes^16^. NFR region lengths are seldom longer than 200 bp (less than 15%) for all species, suggesting that nucleosome-free regions are rarely larger than the span of a single nucleosome.

We aggregated NucleoATAC nucleosome signal around transcription start sites (TSSs) (**Fig. 4d**) to explore species-specific promoter architecture at high resolution. Nucleosome signal proximal to TSSs for both *S. cerevisiae* and humans show a clear depletion at the TSS, with the gap between the +1 and -1 nucleosome slightly larger for the human data. In contrast, this distance is similar to that between adjacent nucleosomes in *S. pombe*; a clear nucleosome free region is not evident, consistent with results from chemical mapping^10^, but not with previous results from MNase that showed a pronounced NFR similar to that observed for *S. cerevisiae*^25^. Others have attributed this discrepancy to the sequence bias of MNase^10^, as the AT rich promoters of *S. pombe* are particularly sensitive to MNase digestion. We also mapped nucleosome occupancy for individual TSSs to determine whether there was heterogeneity in positioning of the +1/-1 nucleosomes that was being masked in the aggregate plot (**Supplementary Fig. 11**). We observe that the pattern observed in aggregate is present for the majority of individual TSS for each organism, although a small fraction of promoters in *S. pombe* do show a larger nucleosome-free and a fraction of promoters in *S. cerevisiae* and human lack a clear nucleosome-free region.

### Dynamic chromatin rearrangements during the osmotic stress response

To demonstrate the ability of NucleoATAC to infer nucleosome positioning and occupancy changes during a dynamic process, we performed ATAC-seq on yeast exposed to osmotic stress (0.6 M increase in the NaCl concentration over 60 minutes). Because osmotic stress induces transient gene expression changes that peak after 15 minutes^26^, we identified promoters with significantly changed accessibility after 15 minutes (FDR < 0.01, fold change > 1.5). In aggregate, the accessibility at these promoters returned closer to steady-state levels during the time-course (**Fig. 5a**), mirroring gene expression changes for these promoters^26^. We analyzed promoters with increasing accessibility during the first 15 minutes of the time course for accompanying shifts and/or depletion of nucleosomes. We identified 3 distinct promoter subtypes characterized by 1) shifts in either the +1 or -1 nucleosomes, 2) decreased occupancy at the -1 nucleosome, and 3) neither of the above patterns (**Fig. 5b**). While shifts in the positioning of +1 or -1 nucleosomes were generally subtle (class 1), nucleosome occupancy declined dramatically for many promoters (class 2), and these promoters also showed greater induction of gene expression. Class 3 promoters, which lacked clear changes in nucleosome positioning despite increased accessibility, showed the least increase in expression **(Fig. 5c)**. These results show that NucleoATAC can reveal transient but biologically relevant chromatin changes during dynamic cellular responses.

**Figure 5.**
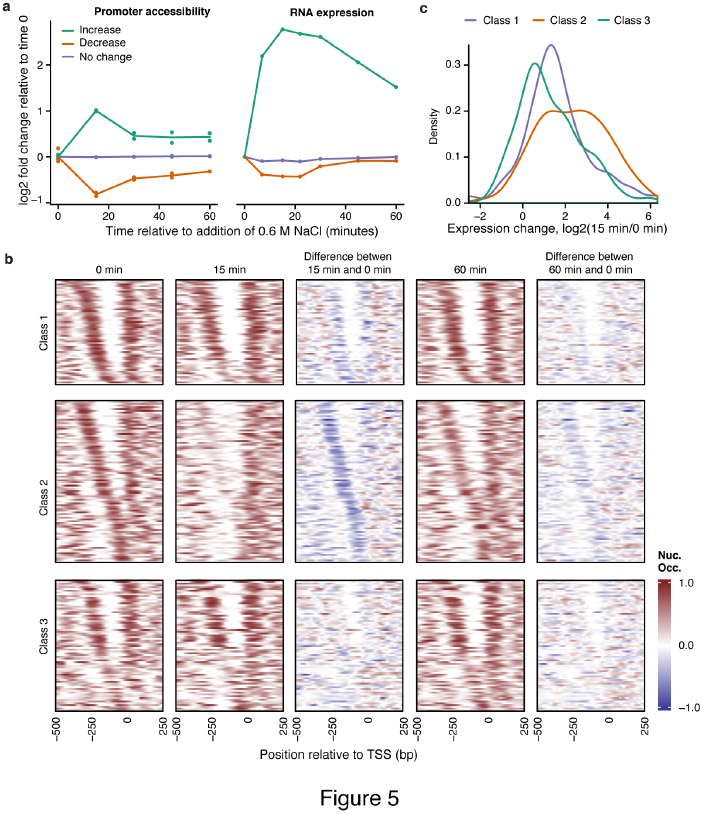
NucleoATAC reveals dynamics of nucleosome positioning and occupancy during osmotic stress response. A) Promoter accessibility (left) and expression (right) changes over the osmotic stress time-course for genes showing an increase in accessibility from 0 minutes to 15 minutes (green), a decrease in accessibility from 0 to 15 minutes (orange), or no significant change in accessibility between 0 and 15 minutes (purple). B) Individual occupancy traces for genes with significantly increased accessibility and belonging to one of three classes in terms of nucleosome positioning/occupancy change: Class 1) shifts in the positioning of the +1 or -1 nucleosomes, Class 2) a decrease in -1 nucleosome occupancy, or Class 3) neither of the patterns for class 1 and 2. C) Distribution of relative expression changes after 15 minutes compared to time 0 for different promoter classes in panel B.

### Transcription factors and nucleosomes compete for binding to DNA

To demonstrate the ability of NucleoATAC to enable detailed investigation of the profiles of human nucleosomes around TFs, we assessed NucleoATAC calls around CTCF motifs. We observe high consistency in the distance of NucleoATAC calls from CTCF binding sites (**Fig. 6a**), consistent with previous observations that CTCF binding sites have highly stereotyped local nucleosome positioning^4,11^. Examining the distance to the nearest nucleosome for other sequence-specific transcription factors, we see that CTCF appears unique in its ability to strongly position flanking nucleosomes, as other TF binding sites often overlap with nucleosomes (**Supplementary Fig. 12**). To explore the relationship between nucleosome occupancy and TF binding more quantitatively, we determined the distribution of nucleosome occupancy for binding sites of 15 sequence-specific TFs (defined as motifs overlapping a ChIP-seq peak) (**Fig. 6b**). For most TFs other than CTCF, a substantial proportion of bound sites have non-zero nucleosome occupancy score, although all show a preference for nucleosome-free DNA. We determined the probability of a TF to be bound based on the motif score and the nucleosome occupancy (**Fig. 6c**). For most analyzed TFs, both nucleosome occupancy and motif strength are predictive of binding, with nucleosome occupancy alone sometimes equally or more predictive than motif score (**Supplementary Figure 13**). This analysis underscores the need for high-resolution nucleosome position and occupancy information to unravel the determinants of TF binding at specific loci.

**Figure 6.**
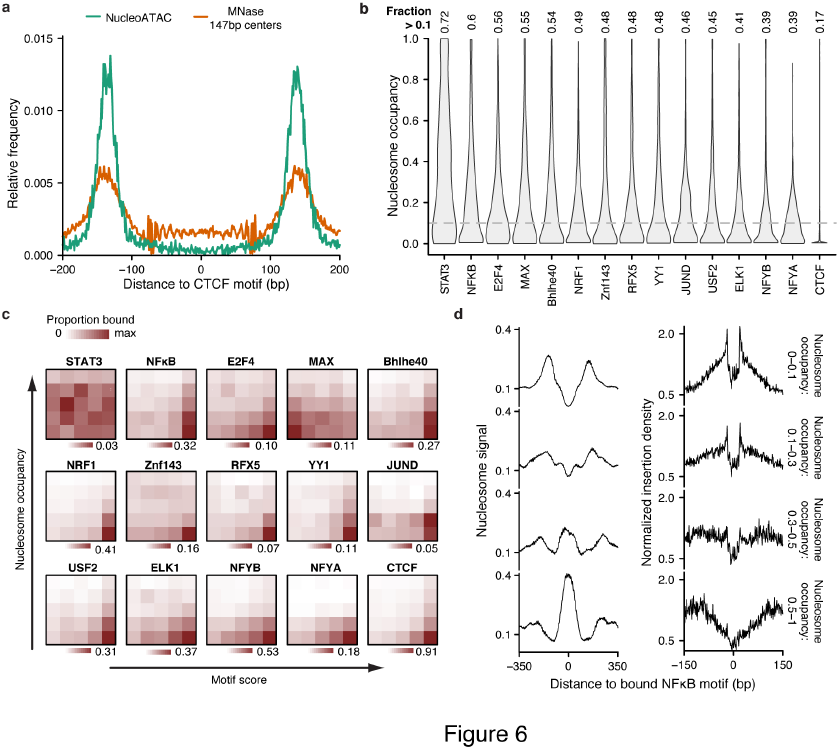
NucleoATAC defines stereotyped TF-nucleosome relationships. A) Nucleosome dyad density relative to CTCF binding site for nucleosomes called with NucleoATAC (teal) or MNase (orange). B) Nucleosome occupancy distributions for sequence-specific TFs. C) Proportion of motifs bound as a function of motif score and nucleosome occupancy. Motif score bins determined by quantiles (0-20^th^ percentile,…, 80^th^-100^th^ percentile); nucleosome occupancy bins determined by occupancy values (0-0.2,…, 0.8-1). D) NucleoATAC nucleosome signal (left) and ATAC-seq insertion profile (right) for NFκB at sites with different nucleosome occupancies. Insertion frequency normalized by sequence bias model.

To further explore the relationship between nucleosomes and TF occupancy, we examined both the nucleosome signal and insertion pattern for NFκB sites with different nucleosome occupancy scores (**Fig. 6d**). Sites with very low nucleosome occupancy exhibited a clear depletion in the nucleosome signal and a clear transcription factor footprint^12^ shown by transposase insertion probabilities (i.e. characterized by a sharp drop in insertions within the motif site). In contrast, high nucleosome occupancy binding sites have a peak in nucleosome signal near the motif and a wide depletion of insertions, indicative of DNA protected by a nucleosome rather than a TF. In addition, sites with high nucleosome occupancy had lower ChIP-seq signal than sites with low or intermediate nucleosome occupancy (**Supplementary Fig. 14**), showing that these sites are indeed less occupied by NFκB in aggregate. Notably, NFκB has been shown to dynamically oscillate between nuclear and cytoplasmic localization^27^; this variablity in localization may provide an explanation for observed intermediate levels of nucleosome occupancy – certain cells have a higher likelihood of being TF occupied vs. nucleosome occupied depending on the nuclear concentration of the TF. All together, these results demonstrate that NucleoATAC may be used to infer dynamic competition between TFs and nucleosomes, with possible applications to understanding the molecular determinants of single-cell regulatory variability.

## Discussion

NucleoATAC utilizes the highly structured 2D fragment size versus midpoint “nucleosome fingerprint” from ATAC-seq chromatin accessibility data to generate high-resolution nucleosome maps within active regulatory elements. These high-resolution maps are highly concordant to those identified by chemical cleavage in yeast and capture the rotational positioning information from nucleosomes in both yeast and humans. As expected, we observe that for *S. cerevisiae* and *S. pombe* WW (W = A or T) and SS (S = G or C) dinucleotides show strong 10-11bp periodicity, but we also observe that all other dinucleotides exhibit considerable periodicity in-phase with the SS dinucleotides. We find human nucleosomes show periodicity in some dinucleotides, but that the extent of periodicity is lower and phasing is less consistent than for both yeast species, suggesting *in vivo* human nucleosome positions are much less constrained by their inherent sequence preference. We also observe a depletion of AT content immediately flanking nucleosomes called by NucleoATAC, in contrast to MNase-based studies that have observed an enrichment of AT content in these regions^4,5^. Together, these data sets validate the hypothesis that human nucleosomes are, in part, positioned by their underlying sequence context *in vivo*, although sequence preferences at the nucleosome boundaries may result from differing sequence biases of the Tn5 and MNase enzymes^7,8^. This observation highlights the need for orthogonal approaches to studying human nucleosome architecture.

Most methods for determining nucleosome positions measure nucleosome depletion only indirectly through a lack of (unnormalized) signal. In contrast, ATAC-seq simultaneously assays nucleosome depletion (through the presence of short fragments) and nucleosome positioning (from longer fragments). Combining measurements of chromatin accessibility, nucleosome positioning, and nucleosome occupancy allows an integrative analysis of chromatin architectural changes, as demonstrated by the observation of two classes of promoter changes during the osmotic stress response in *S. cerevisiae*. In contrast to MNase-based assays that found nucleosome position/occupancy were not predictive of expression dynamics^28–30^, our integrated analysis shows that promoters with decreasing occupancy of the -1 nucleosome showed stronger increases in expression compared to other dynamic changes in promoter architecture. These results suggest that NucleoATAC can reveal distinct mechanisms causing promoter accessibility that lead to different magnitudes of gene expression change.

High resolution nucleosome calls and occupancy tracks also enabled investigation of the effects of nucleosome occupancy on TF binding within regulatory regions. For many factors, nucleosome occupancy is predictive of TF binding, suggesting that incorporation of nucleosome positioning and occupancy information for these TFs could improve algorithms for inferring TF binding using ATAC-seq^31,32^. Some factors show intermediate nucleosome occupancy at a substantial fraction of binding sites, suggesting cell-to-cell heterogeneity in TF occupancy. This heterogeneity may be linked to oscillation in nuclear localization of TFs;^33^ examining which sites show partial occupancy may reveal new insight into how TF pulsing dynamics influence binding and gene regulation.

NucleoATAC provides a powerful new framework for analyzing ATAC-seq or other paired-end functional genomics data. By using a two-dimensional fragment size versus midpoint representation of ATAC-Seq, NucleoATAC is built on the understanding that different fragment lengths provide unique information content. Our 2-D signal processing approach can likely be extended by applying additional methodologies from the image analysis field to the 2-dimensional fragment sizes versus midpoint “image” of ATAC-seq data, opening exciting possibilities for future applications for calling genomic features other than nucleosomes.

ATAC-seq coupled with NucleoATAC allows for the interrogation of high-resolution nucleosome positions in regulatory regions from limited cellular populations, allowing rapid, cost-effective and high-resolution nucleosome inference. We believe future efforts will include fine mapping of chromatin structure in rare developmental and disease cellular populations, providing a detailed understanding of the molecular determinants of chromatin structure across dynamic cellular processes in human cells.

## Materials and Methods

### Yeast strains, growth, and library preparation

Four populations of *S. cerevisiae* strain GSY147^34^ were used, with three of the four populations derived from the first after being grown on plates for 200 generations in conditions to minimize selective pressures. Replicates were grown to mid-log phase in YPD, then 5 million cells per library were harvested. The cells were washed twice in Sorbitol buffer (1.4 M Sorbitol, 40 mM HEPES-KOH pH 7.5, 0.5 mM MgCl_2_) then incubated for 30 min at 30°C and 300 rpm shaking with 0.5 mg/mL 100T zymolyase. The cells were washed twice in Sorbitol Buffer, then incubated with 2.5 μl of Nextera Transposase in 47.5 μl of 1x TD buffer at 37°C for 30 minutes. PCR was performed as described in Buenrostro et al. (2015)^35^. The libraries were sequenced with 50bp paired-end reads on an Illumina HiSeq.

For the time course analysis of osmotic stress, *S. cerevisiae* strain BY4741 was grown in YPD until 0, 15, 30, 45, or 60 minutes before harvesting, at which point a 5M NaCl solution in water was added for a final concentration of 0.6 M NaCl. Two replicates were used for each time point, except for time 0 for which there were four replicates. For all time points other than two of the replicates for time 0, Sorbitol Buffer washes and incubations prior to transposition were carried out in buffer with 0.6M NaCl. To accelerate the time between harvesting the cells and performing the transposition, the protocol was modified as follows: Cells were washed only once in Sorbitol buffer with 10mM DTT before being incubated for 5 minutes at 30°C and shaking at 300 rpm with 0.5 mg/mL 100T zymolyase in Sorbitol Buffer with 10mM DTT. Cells were then washed once with Sorbitol buffer before being incubated with 2.5 μl of Nextera Transposase in 47.5 μl of 1x TD buffer at 37°C and shaking at 300 rpm for 15 minutes. PCR was performed as described in Buenrostro et al. (2015)^35^. Libraries were sequenced with 76 bp paired-end reads on an Illumina NextSeq 500.

*S. pombe* strain 972 h-was grown to mid-log phase in YES media, and 1, 5, or 20 million cells were washed twice in Sorbitol Buffer + 10 mm BME, incubated in Sorbitol Buffer + 2mM BME for 30 minutes, and the incubated in Sorbitol buffer with 0.25 mg/mL 100T Zymolyase for 5 or 23 minutes, and then washed twice in Sorbitol Buffer before incubated in 10 μl of 1x TD buffer and 0.5 μl of Nextera Transposase at 37°C for 30 minutes. PCR was performed as described in Buenrostro et al. (2015)^35^. Libraries were sequenced with 76 bp paired-end reads on an Illumina MiSeq.

### Sequencing alignment and peak calling

Bowtie2 was used to align *S. cerevisiae* reads to the sacCer3 genome (April 2011 Release from *Saccharomyces* Genome Database^36^), *S. pombe* reads to the ASM294v2.21 genome, and GM12878 reads to the hg19 genome. Duplicates were removed for each individual library using Picard tools. Reads with mapping quality below Q30 as well as improperly paired reads were removed from subsequent analysis. For the 11 *S. cerevisiae* samples that were not part of the osmotic time course, replicates across all biological samples were merged for all downstream analysis (unless otherwise specified). The 5 *S. pombe* replicates, 4 GM12878, and the 2 *S. cerevisiae* samples for each time point in the osmotic stress time-course were also merged for nucleosome analysis.

For the two yeast species, highly mappable regions were determined using custom 35 bp alignability tracks; only these mappable regions were used for subsequent analysis. For the GM data, open chromatin regions were called using MACS2 with the broad flag. Peaks were filtered based on mappability using the Duke 35 bp alignability track for hg19. All subsequent analysis of GM data was performed only for these mappable open chromatin regions.

### Insertion position and fragment size determination

The start of sequencing reads generated from ATAC-seq are offset from the center of the Tn5 binding site by 4 bp^11^. Thus ATAC-seq insertions were defined as single base-pair sites 4 bp from the ends of sequencing fragments. Similarly, fragment size was defined as the size of the sequenced fragment minus 8 base pairs so that fragment size represents the distance between the centers of two Tn5 binding sites.

### Occupancy determination

We sought to model the fragment size distribution as a mixture of nucleosome-free fragments and nucleosome-associated fragments in a way that captured the highly structured, non-parametric nature of the nucleosomal distribution. Because fragments less than 115 bp very likely arise from the nucleosome-free distribution, we parameterized the fragment size distribution below that size-cutoff as an exponential distribution, which provided a good fit to this region of the distribution. This fit distribution was used to extrapolate the nucleosome-free fragment distribution for sizes larger than 115 bp. The subtracted difference between the extrapolated nucleosome-free model and the observed fragment distribution was used as the nucleosome-associated fragment distribution. Nucleosome occupancy at a particular site in an open chromatin region was determined by computing the fragment size distribution within a Gaussian window centered at that position with standard deviation of 25 bp, then modeling this local distribution as a mixture of the nucleosomal and nucleosome-free insert size distributions: *P(i)* = *α* * *P*_*nucleosomal*_*(i)* + (1−*α*) * *P*_*nucleosome-free*_*(i)*, where α represents the fraction of fragments arising from the nucleosomal distribution. The maximum likelihood estimate for α defined the occupancy, which ranges between 0 and 1.

### V-plot normalization

The yeast V-plot used for cross-correlation was generated by aggregating reads around dyad calls from chemical mapping that met two criteria: (1) They had an NCP/noise ratio (positioning metric defined by Brogaard et al.^9^) in the top 20% of calls, and (2) they had nucleosomal occupancy (determined as described above) of greater than 0.5. The portion of the V-plot representing fragments of sizes between 105 and 250 bp with fragment centers within 60 bp of the dyad position was normalized to match the nucleosomal fragment size distribution of the sample being analyzed. For this normalization, we used the initial mixture model for the fragment size distribution to determine a refined nucleosome-associated fragment distribution. Peaks in the nucleosome occupancy track—as determined from the initial model—were identified as candidate, low-resolution nucleosome positions. The fragment size distribution for fragments centered within 60 bp of these peak positions was then used as the nucleosomal fragment size for the v-plot normalization. Each row in the V-plot corresponds to a specific fragment size; the elements in a particular row were all scaled so that the sum of the row would match the frequency of that fragment size in the nucleosome-associated fragment size distribution. The V-plot was also symmetrized across the vertical axis and smoothed slightly with a Gaussian filter with standard deviation of 1 bp.

### Nucleosome signal track and background subtraction

This V-plot was cross-correlated against matrices defining the fragment center and size information for a genomic region, such that the cross-correlation signal at position x along the genome is given by *Single(x) = F·V* where *F* is the matrix of fragment center and size information for fragments of size 105 to 250bp with centers between *x* - 60 and *x* + 60 and V is the V-plot matrix. This raw signal is then normalized using a backround signal that is intended to represent the expected signal from the cross-correlation given 1) the number of fragments observed and 2) the Tn5 sequence preference. The background signal at position x is defined as *Background(x)* = *B · V \**Σ*F*, where *B* represents a matrix with relative probabilities of generating fragments of different sizes and center positions such that Σ*B* = 1. The scaling factor Σ*F*, the sum of all reads in the signal matrix, ensures that the background signal represents the expected signal given the observed number of fragments. To determine *B*, the probability of observing individual insertion sites was first modeled as follows. Tn5 has a sequence preference across about 21 bp that it contacts^11^, therefore we developed a Position Weight Matrix (PWM) for sequence content +/- 10 bp from Tn5 insertion points in ATAC-seq performed on genomic DNA. Relative probabilities are calculated for each genomic position using this PWM, and then this 1D sequence preference is used to calculate the relative probability of observing particular ATAC-seq fragments (which require two Tn5 insertions) by multiplying the probabilities of the two insertions needed for that fragment with the probability of observing a fragment of that size (determined from the fragment size distribution). The normalized nucleosome signal is given by subtracting this background signal from the cross-correlation signal: *Normalised Signal(x) = F · V − B · V \** Σ*F*.

### Calling dyad positions

The normalized nucleosome signal tends to be highly periodic with many local maxima. To robustly identify maxima representing potential nucleosome dyad positions while still preserving the rotational positioning information in the periodic signal, the normalized signal is smoothed using a Gaussian window of 25 bp and local maxima are found in the sum of this smoothed signal and the original normalized signal. These local maxima are considered candidate nucleosome positions. To define a non-redundant map of nucleosome positions, a greedy algorithm is employed in which the candidate nucleosome position with the highest signal is chosen to be included in the map, then the next highest peak not within 120 bp of any position in the map already, until no peaks remain that are not within 120 bp of the non-redundant set.

For each dyad, a Z-score is determined by calculating *Z = (F · V − B · V \** Σ *F)/var (Background)* where the variance of the background signal based on the bias model is 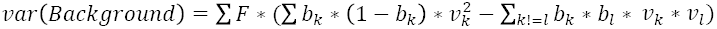, with *b*_*k*_ and *b*_*l*_ as individual elements of matrix B and *v*_*k*_ and *v*_*l*_ the corresponding elements of matrix V. A log likelihood ratio is also determined by calculating the likelihood of the data given that the data arises from the V-plot pattern multiplied by the local bias pattern and the likelihood of the data arising purely from the local bias pattern. For all analyses, only calls with Z-scores greater than 3, log likelihood ratios greater than 0, and occupancies greater than 0.1 were considered. These cutoffs were chosen based on three factors: 1) Analysis of ATAC insertion profile and V-plot around calls, 2) Concordance between calls and chemical mapping, and 3) Concordance between calls for individual replicates. As can be seen in Supplementary Figures 6-8, increasing the stringency of thresholds leads to more consistent calls that are more concordant with chemical mapping. The choice of thresholds for different applications should be based on the desired balance between more comprehensive calls and higher confidence calls.

### Dinucleotide pattern analysis

For comparison of dinucleotide frequencies between species, a higher confidence threshold was used—only calls with log likelihood ratios greater than or equal to 5 were used (threshold chosen based on analysis of random down-samples and individual replicates, as in **Supplementary Fig. 6-7**). For both correlation and power spectrum density analysis, dinucleotide frequencies between 19 and 60 bp from the dyad calls (averaged across both sides of the dyad as the calls were not inherently stranded) were normalized by division with the mean frequency of the dinucleotide in that window. The power spectrum density at frequency 1/10.5 was calculated as the square of the fast Fourier transform at that frequency.

### Osmotic stress time-course analysis

For differential accessibility analysis between time points, we compared the number of insertions in the 300 bp window upstream of the TSS between time points. As variation in the degree of enrichment of fragments within open chromatin regions can affect differential accessibility measurements between ATAC-seq samples, counts were normalized as follows: Quantile normalization was applied to the counts for promoters with genes showing no expression change as determined by a previous study^26^ (Raw counts for these genes were highly correlated). A lowess curve was then fitted to the quantile-normalized counts versus raw counts for each sample to map the smooth transform to be applied to the raw counts in that sample. This transform was then applied to all genes. The natural log of the difference between the normalized and raw counts was used as an offset in a GLM model regressing observed counts on the salt induction time point using the R package edgeR^37^. A likelihood ratio test was performed to identify any gene that changed significantly over any interval during the time-course. Genes with a significant change over the time-course at an FDR of 0.01 and with a 1.5 fold change in normalized accessibility between 15 minutes and 0 minutes were selected as having increased or decreased significantly in the first 15 minutes.

For genes with increased accessibility, changes in nucleosome patterning were determined as follows: The +1 and -1 nucleosome positions relative to TSS were determined using NucleoATAC signal and occupancy: 1) Genes with shifts of the -1 nucleosome 8 to 50 bp upstream and/or shifts of the +1 nucleosome 8 to 50 bp downstream with decreases in occupancy for the -1 nucleosome less than 0.1 were assigned as showing “+1/-1 nucleosome shifts.” 2) For the remaining genes, if the occupancy at the position of the -1 nucleosome at time 0 decreased by more than 0.1 between 0 and 15 minutes, the gene was assigned as undergoing “-1 nucleosome depletion”. 3) The remaining genes were classified as “other”.

### Downloaded datasets and annotations

Chemical mapping data for *S. cerevisiae* were obtained from Supplementary Table 2 from Brogaard et al. (2012)^9^ and lifted over to the sacCer3 genome. Chemical mapping for *S. pombe* were obtained from Supplementary Dataset 01 from Moyle-Hermann et al. (2013)^10^. MNase nucleosomes calls labeled as “MNase 1” were obtained from http://compbio.cs.huji.ac.il/NucPosition/TemplateFiltering/Nucleosome_calls_files/BY_10uL_normal_digestion.tab^20^ and lifted over to sacCer3. MNase calls labeled as “MNase 2” were obtained by applying DANPOS2^17^ to the MNase sequencing data obtained from SRR094649 and SRR094650^21^. Human (GM) MNase fragment center positions mapped to hg19 were obtained from the Pritchard lab by request.

For *S. cerevisiae*, TSS were determined using median UTR lengths from TIF-Seq^38^ and gene annotations from the *Saccharomyces* Genome Database^36^. *For S. pombe*, TSS were obtained from Supplementary Table 2 from Lantermann et al. (2012)^25^. For human, TSS were defined by CAGE signal from the ENCODE project^39^; for each transcript, only the TSS with maximum CAGE signal was used. For Fig. 6, uniformly processed ENCODE/SYDH ChIP-seq datasets were downloaded from the UCSC ENCODE data repository (http://hgdownload.cse.ucsc.edu/goldenPath/hg19/encodeDCC/wgEncodeAwgTfbsUniform/). ChIP-Seq peaks were intersected with motif occurrences called using FIMO^40^ and the JASPAR database.

### Accession numbers and software availability

The raw data for *S. cerevisiae* and *S. pombe* as well as nucleosome positions and signal tracks for all three species analyzed have been deposited at GEO accession number GSE66386. Raw data for the GM12878 cell line are available at GEO accession number GSM1155960.

### Code availability

NucleoATAC source code is freely available as a python package at https://github.com/GreenleafLab/NucleoATAC.

## Acknowledgements

ANS, JDB, and SDK are supported by NSF GRFP; ANS and JDB also acknowledge the NIH training grant T32HG000044 for support. This work was supported by National Institutes of Health (NIH) P50HG007735 and U19AI057266, and the Rita Allen Foundation (to W.J.G.) The Stanford Genetics and Developmental Biology entering PhD students in 2014 assisted in the osmotic stress time-course experiment as part of the 1^st^ Year Training Camp. We also thank members of the Greenleaf and Chang labs for useful discussions, Anshul Kundaje for feedback on the manuscript, Paul Giresi for input on the yeast ATAC-seq protocol, and the Pritchard lab for 147 bp MNase centers mapped to hg19.

## Author Contributions

All authors contributed to the design of the experiments. SKD, KS, JDB, and ANS performed the experiments. ANS, JDB, and WJG conceived the analysis framework. ANS wrote the NucleoATAC software and performed the data analysis. ANS and WJG wrote the paper with feedback from all authors.

## Supplementary Materials

Supplementary Figures 1 to 14

Supplementary Software

**Accession Numbers**

GSE66386

GSM1155960

